# A role for YY1 in sex-biased transcription revealed through X-linked promoter activity and allelic binding analyses

**DOI:** 10.1101/044172

**Authors:** Chih-yu Chen, Wenqiang Shi, Allison M. Matthews, Yifeng Li, David J. Arenillas, Anthony Mathelier, Masayoshi Itoh, Hideya Kawaji, Timo Lassmann, FANTOM Consortium, Yoshihide Hayashizaki, Piero Carninci, Alistair R. R. Forrest, Carolyn J. Brown, Wyeth W. Wasserman

**Affiliations:** Centre for Molecular Medicine and Therapeutics, Child and Family Research Institute, University of British Columbia, Vancouver, British Columbia, Canada; Graduate Program in Bioinformatics, University of British Columbia, Vancouver,British Columbia, Canada; RIKEN Omics Science Center, Yokohama, Japan; RIKEN Center for Life Science Technologies, Division of Genomic Technologies,Yokohama, Japan; RIKEN Preventive Medicine and Diagnosis Innovation Program, Wako, Saitama, Japan; Department of Medical Genetics, University of British Columbia, Vancouver, British Columbia, Canada

**Author notes:** Corresponding authors (WWW); (CJB).

## Abstract

Sex differences in susceptibility and progression have been reported in numerous diseases. Female cells have two copies of the X chromosome with X-chromosome inactivation imparting mono-allelic gene silencing for dosage compensation. However, a subset of genes, named escapees, escape silencing and are transcribed bi-allelically resulting in sexual dimorphism. Here we conducted analyses of the sexes using human datasets to gain perspectives in such regulation. We identified transcription start sites of escapees (escTSSs) based on higher transcription levels in female cells using FANTOM5 CAGE data. Significant over-representations of YY1 transcription factor binding motif and ChIP-seq peaks around escTSSs highlighted its positive association with escapees. Furthermore, YY1 occupancy is significantly biased towards the inactive X (Xi) at long non-coding RNA loci that are frequent contacts of Xi-specific superloops. Our study elucidated the importance of YY1 on transcriptional activity on Xi in general through sequence-specific binding, and its involvement at superloop anchors.

## Background

Sex disparities in disease progression and susceptibility for many diseases, including cancer, autism, cardiac and autoimmune disorders, have been long known. Such discrepancies likely result from a combination of the sex chromosomes, sex hormones, and environmental factors. Due to such sexual dimorphism, there has been a recent push in policy to balance sex in cell and animal studies by NIH ^1^. The key genetic differences between the sexes are the sex chromosomes, with mammalian females being XX and males XY. With the exception of the pseudoautosomal regions (PAR) shared with the Y-chromosome, X-linked genes are present in 2 copies in females and only 1 in males. X-chromosome inactivation (XCI) silences one copy of the X chromosome (chrX) in female cells in order to compensate for dosage between the sexes. Up-regulation of *XIST*, a long non-coding RNA (lncRNA), is known to be responsible for initiation of XCI, and this process has been reported to be mediated through recruitment of factors such as polycomb repressive complex 2, leading to tri-methylation of histone H3 at lysine 27 (H3K27me3) and silencing of the inactive X chromosome (Xi) ^2^. XCI involves the establishment of a peripheral nuclear architecture, which includes association with the lncRNA, *FIRRE*^3^, anchoring the Xi near the nucleolus to preserve H3K27me3 and the silencing state^4^ DNA methylation (DNAm) is recruited to promoters, providing maintenance of the inactive state. As a result, the majority of chrX genes outside of PAR regions are subject to XCI, and are transcribed mono-allelically from the active X (Xa) in female somatic cells. A small subset of chrX genes including *XIST*, known as escapees, escape from XCI and are transcribed on the Xi. These escapees are therefore bi-allelically transcribed with the exception of *XIST*, which is solely transcribed from the Xi. Binding of the Ying-Yang 1 (YY1) transcription factor (TF) to *XIST* RNA and DNA contributes to *XIST* transcription ^5-7^. *XIST* and *FIRRE* are among the four long non-coding RNAs (lncRNAs) previously found at frequently interacting regions of Xi-specific superloops in the GM12878 cell line ^8^. Rao *et al*. reported tandem CTCF motifs at three (*FIRRE*, *DXZ4*, *LOC550643*) of the four lncRNAs, and suggested a role for chromatin looping through CTCF and RAD21 in shaping the chromatin structure of Xi. At the *FIRRE* locus, Yang *et al*. showed differential occupancy by CTCF and YY1 using ENCODE ChIP-seq peaks from male and female cells ^4^.

Investigation of differences between the sexes has been tackled both through direct comparisons of male and female data and/or a focused study of XCI. Direct sex comparisons have identified male:female differences in gene expression, DNAm, accessible chromatin, and TF binding levels. At the gene expression level, microarray and RNA-seq platforms have been used to examine differential gene expression between the sexes. The Genotype-Tissue Expression (GTEx) pilot study reported genes with differential expression between male and female samples in 43 tissues using RNA-seq data ^9^. Hall *et al*. integrated RNA-seq, DNAm, and microRNA datasets in islet cells for a sex comparison to identify genes associated with insulin secretion, revealing the potential molecular mechanism for phenotypic sex differences in this tissue ^10^. At the DNAm level, we previously focused on chrX to compare between sexes in 27 tissues using Illumina 450K arrays, from which we identified escapees and subject genes across cell types ^11^. The assay of transposase accessible chromatin with sequencing (ATAC-seq) maps chromatin accessibility in a given cell population. A sex comparison on chrX by Qu *et al*. using ATAC-seq data in T cells revealed the highest female to male ratio of chromatin accessibility at *XIST* and *FIRRE*, followed by escapees^12^.

Allelic expression or allelic binding approaches are powerful in delineating activities from either Xi or Xa, although they require cells with XCI of one chrX favored over the other (skewed XCI) as well as the genotype information of the cells. These studies in general are limited by the availability of known heterozygous sites within the particular cells for allelic comparison. Experiments using F1 mouse cells containing large numbers of known heterozygous sites have therefore been favored. In human lymphoblastoid cell lines, allelic expression and histone modifications, measured using Illumina BeadChip genotyping arrays, showed the majority of genes to be subject to XCI, and a correspondence between expression and histone modifications ^13^. Allelic RNA-seq and ChIP-seq for RNA polymerase II and various TFs have previously been conducted on the lymphoblastoid GM12878 cell line known to have skewed XCI ^14-16^. Reddy *et al*. reported that TFs are predominately bound on the Xa in GM12878, and allelic binding of RNA polymerase II are non-differential between Xa and Xi at escapees ^16^.

The recent generation of high throughput datasets offers new opportunities to examine X-linked gene expression. The cap analysis of gene expression (CAGE) datasets, which measure abundance of the 5’ end of transcribed RNAs, generated by the FANTOM5 consortium have facilitated the identification of transcription start sites (TSSs) and their levels of transcription in over 800 samples of non-treated human cells^17^. The strength of CAGE datasets compared to RNA-seq lies in the capability of pinpointing TSSs of gene promoters as well as enhancers ^17,18^. For the ease of terminology, we refer to the reported robust CAGE peaks as TSSs in the rest of the text. In addition, The Cancer Genome Atlas (TCGA) offers a rich source of DNAm datasets of both sexes in specific cancer types from the epigenetic perspective ^19^, while over 680 TF ChIP-seq datasets generated by the ENCODE project provide information on TF binding events in multiple cell types ^20^.

Here we report a comprehensive investigation of X-chromosome gene activity between the sexes, focused on escapees through direct sex comparisons of expression, DNAm, and TF binding levels, further supported with evidence of allelic binding on chrX in the female GM12878 lymphoblastoid cell line. While XCI has previously been reported to occur at a domain level ^21,22^, we hypothesize that there exists a common regulatory mechanism that facilitates escape from XCI. To our knowledge, this is the first study conducting large-scale analyses on TF ChIP-seq data comparing between sexes and between Xi and Xa occupancies to identify potential regulator(s) associated with escape from XCI. The work is part of the FANTOM5 project ^17,23^. Data downloads, genomic tools and co-published manuscripts are summarized at http://fantom.gsc.riken.jp/5/.

## Results

### Sex classification of the FANTOM5 samples using CAGE data

To understand how well the transcriptional activity of TSSs can distinguish sexes and to confirm reliable labeling of male and female samples, we built a sex classifier through a Random Forest approach ^24^ using the FANTOM5 CAGE dataset. We trained on 530 samples of known sexes using the transcriptional levels of 5071 TSSs in the non-pseudoautosomal region of the X (X-non-PAR) as features. Through performance assessment, the classifier performed better than a classifier based on TSSs assigned to *XIST* alone. Namely, the out of bag error rate and balanced accuracy were 6.51% and 0.908 for the X-non-PAR classifier and 11.55% and 0.825 for the *XIST* classifier. Hence, we used results from the X-non-PAR classifier in our following analyses. The first two dimensions of multi-dimensional scaling of the sample proximity values from the Random Forest classifier indicated the presence of outlying samples (Fig 1A; S1 Table). The *XIST* TSSs of these outlying samples were differently expressed from their original sex labels. Of the 26 samples called female but classified male, there were 2 stem cells and 14 cancer lines, in which the *XIST* expression were zero to minimal. The latter group is consistent with reports of certain cancer cells losing the Xi ^25^. Of the 10 samples labeled male but classified female, there were 3 testes-related tumor cell lines with strong *XIST* expression in agreement with a previous report of testicular germ cell tumors expressing *XIST* ^26^. The remaining samples could reflect sex chromosome aneuploidy or sample mislabeling. For the latter we noted at least one case where the labeled sex differed between technical replicates from the same source. Overall, the results suggested that we can classify sexes accurately using transcriptional activity of TSSs in X-non-PAR regions. Thus we constructed our sex classifier using X-non-PAR TSSs training on all labeled samples, and predicted a total of 309 female and 426 male samples to be used in the subsequent section with the exclusion of outlier and mixed samples (S1 Table).

**Figure 1:**
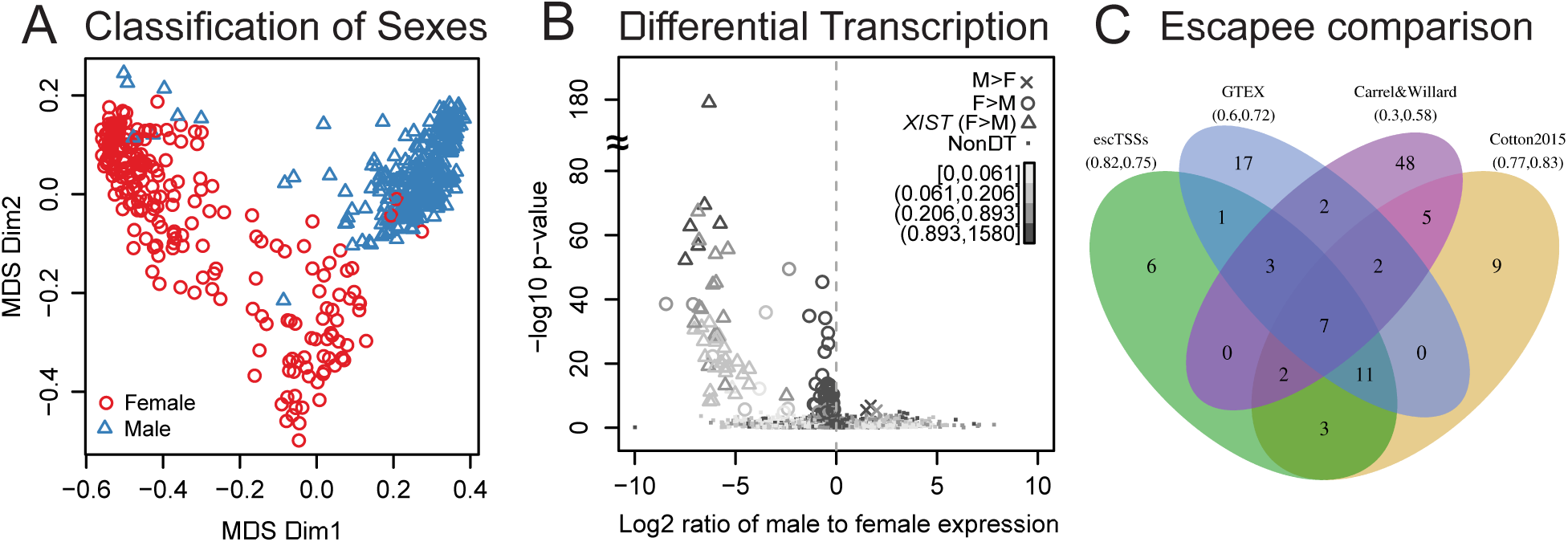
Classification and differential expression analysis of sexes using FANTOM5 CAGE datasets. (**A**) Plot showing distance between FANTOM5 CAGE samples in the first 2 dimensions generated from multi-dimensional scaling of the proximity matrix from the Random Forest sex classifier. Each point represents a FANTOM5 sample with its labeled sex: male (blue triangle) or female (red circle). (**B**) Scatter plot with log2 ratio of the mean expression levels comparing male to female (with a constant of the 5^th^ percentile expression added to avoid denominators of 0) on the x-axis, and ‐log10 transformation of raw p-values from the differential transcription analysis between sexes on the y-axis. Each point represents a TSS in non-PAR region of chrX, and TSSs with significantly higher expression in female (escTSSs) and male cells are denoted with circles and crosses (Bonferroni-corrected p-value ≤ 0.05), with small dots for non-differentially transcribed TSSs. The escTSSs nearest to the *XIST* gene are denoted using open triangles. The grey-scale gradient represents the average expression across all samples in quartiles. The vertical dashed line represents a log2 ratio of 0, where there is no difference between sexes. (**C**) Venn diagram depicting the overlapping sets of escapees from three published studies with those identified in this report. The numbers within the Venn diagram represent the overlaps between sets, and the numbers in bracket under each list name are precision and recall values where genes reported in more than one list are taken to constitute true escapees.

### Differentially transcribed chrX TSSs with higher expression in female reflect escapees

Given that there were disproportionately large numbers of samples of one sex for multiple cell types, we incorporated cell categories from ontologies to group multiple closely related cell types to minimize bias from cell-type specific transcription. Using the sex labels from our classifier, we assessed the differential transcription between sexes using a linear regression model for each TSS, incorporating the cell categories as covariates (see Methods for details). Parallel analysis was conducted on both the X-non-PAR region and autosomes (S2 Table). Here we focused on chrX, with some information from the autosome analysis presented in S1 File. Of the 103 TSSs on chrX that were significantly differentially transcribed between sexes, 3 of the 5 TSSs assigned to the *ARHGAP4* gene had higher expression in male cells, while 100 TSSs (referred to as ‘escTSSs’) corresponded to 33 unique genes with higher expression in female cells (Fig 1B). Forty-eight of the 100 escTSSs were associated to the *XIST* gene, and exhibited much stronger differences between sexes than others (Fig 1B).

Given the higher expression in female cells, we expected the majority of our predicted escTSSs to escape from XCI (S1 Fig A). We compared the gene name of our escTSSs to previously identified escapees from the literature, in which different data and techniques were used: RNA-seq (‘GTEx’) ^9^, rodent/human somatic cell hybrids (‘Carrel&Willard’) ^27^, and Illumina 450k DNAm array (‘Cotton2015’) ^11^. For our comparison, the set of escapees identified by at least two approaches were assumed to be true escapees. Our approach obtained the highest proportion of overlap with the escapees (precision=0.82), and ranked second after ‘Cotton2015’ for the proportion of the escapees called (recall=0.75; Fig 1C). The top 4 of the 6 genes from our list that were not captured by previous literature were non-coding RNAs: *RP13-348B13.2*, *U2*, *MIR4767*, and *RNF19BPX*. Out of the 9 genes not identified (false negatives), two genes, *GYG2* and *STS*, were reported in three other approaches. We note a connection between a pair of entries in the two categories, as the *MIR4767*gene TSS overlaps with a *STS* transcript according to the UCSC gene annotation of the hg19 assembly. Summarizing the key point, the escTSSs identified from differential transcription of the sexes predicted escapees with the highest precision.

### DNA methylation similarity between sexes on chrX showed strong agreement with bi-allelically transcribed escapees

We have previously used DNAm data to identify genes that were subject to or escaping from XCI ^11^. Here we used an independent public DNAm dataset from TCGA ^28^ to examine whether the differential transcription we identified was reflected at DNAm level between the sexes. On autosomes, we expected differential DNAm between the sexes to correspond to differential transcription. On chrX, however, the overall methylation level was reported to be higher in female than in male due to the nature of the Xi ^11^, but we expected similar DNAm levels between sexes at bi-allelically transcribed escTSSs (S1 Fig B). For ease of reference, we refer to bi-allelically transcribed escTSSs as ‘bi-escTSSs’ for the rest of the text. DNAm of probes near TSSs of *XIST*, *ZFX* (a bi-allelically transcribed escapee), and *UPF3B* (a subject gene) are shown as examples in Fig 2A using data in female and male urothelial bladder cancer samples from TCGA.

**Figure 2:**
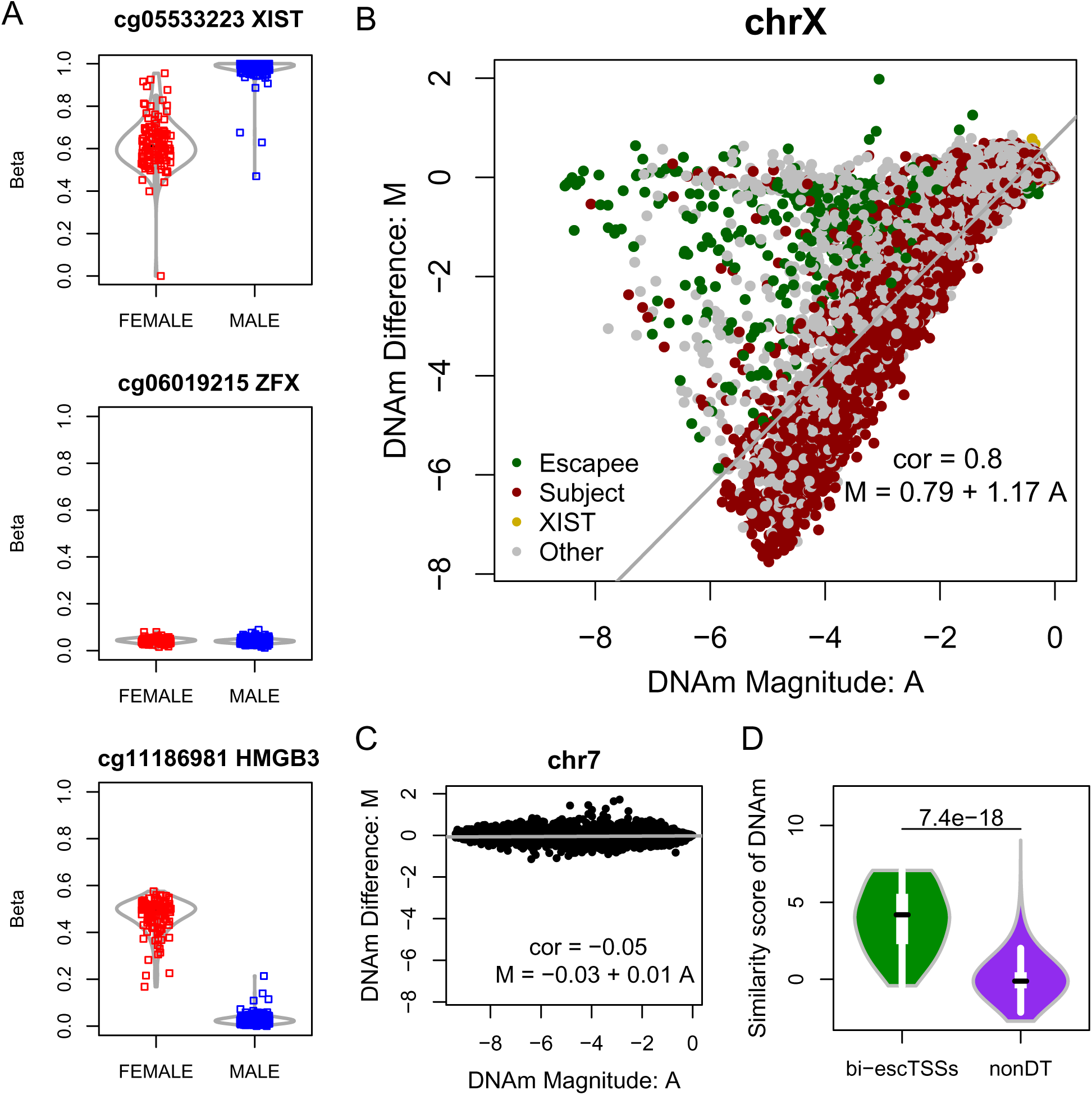
DNAm comparison between sexes on chrX in urothelial bladder cancer (BLCA) samples from TCGA. (**A**) DNA methylation status for positions (i.e. probes from the Illumina 450k array) near TSSs in both sexes from BLCA samples, where the *β* values (Y-axis) range from 0 (unmethylated) to 1 (fully methylated). The three TSSs are most proximal to the following genes (from top to bottom): *XIST*, an escapee (*ZFX*) and a subject gene (*HMGB3*). Each square represents a sample for the BLCA dataset. Red or blue color represents a female or male sample, respectively. Each violin plot in gray lines shows the distribution of beta values for each sex at each probe. Plots (**B**) and (**C**) show MA plots for chrX probes and autosomal probes on chr7 between sexes, respectively. Each dot represents a probe from the array. M (difference) on y-axis is the logged differential methylation value between sexes, and A (magnitude) on x-axis is the logged average methylation value (as indicated in Methods). The fitted robust regression line is represented in gray, with the corresponding function and correlation reported. Green and red colors in plot (**B**) represent probes nearest to escapees and subject genes previously reported in Cotton *et al*. 2015. Gold and gray colors represent probes nearest to *XIST* and genes not in either three categories. (**D**) Violin plots showing the distributions of DNA methylation similarity scores between sexes for probes within 50bps of escTSSs and non-differentially transcribed (nonDT) TSSs on chrX. The similarity score of DNA methylation on y-axis is the residual of M as a function of A on chrX. Only TSSs with at least one probe within 50bps were plotted, and for those TSSs within 50bps of multiple probes, the average similarity scores of probes were obtained. The p-value from the Wilcoxon test is reported above the violin plots.

To show the differences in DNAm between sexes on chrX and autosomes, we generated microarray-inspired MA plots to compare the relationship between log2 ratios of DNAm between male and female samples (i.e. the difference: M) versus log2 average of DNAm between the sexes (i.e. the magnitude: A) for probes in chrX and chr7 (Figs 2B and 2C; see methods for details). We discovered a drastically stronger linear correlation between difference and magnitude on chrX (ρ=0.8) but not on autosomes (chr7: ρ=-0.05). On autosomes, probes that deviate from the zero difference values were differentially methylated between sexes; whereas on chrX, probes that deviate from the fitted regression line, with the log beta ratio closer to 0, had similar DNAm levels between sexes. Hence, we identified the chrX probes with similar DNAm levels between sexes by computing the residual of the regression model as the similarity score, which also gives greater weights to probes with lower DNAm levels. Indeed, the DNAm similarity scores from probes within 50 base pairs (bps) of TSSs were found to be significantly higher for bi-escTSSs than non-differentially transcribed TSSs (nonDT; p-value of Wilcoxon test: 7.4*10^‒18^; Fig 2D). We observed similar results using three other cancer datasets (S2 Fig). Taken together, the results showed that higher DNAm similarity between sexes on chrX reflected TSSs of escapees, and further supported the bi-allelic activities of the bi-escTSSs we predicted.

### YY1 binding motif over-representation around escTSSs

Given that escTSSs (both bi-escTSSs as well as *XIST*) are transcribed on the Xi, we began testing our hypothesis that common regulator(s) and mechanism exist to facilitate the escape from XCI. We first probed the regulation of the escTSSs on chrX at the sequence level through enrichment testing of JASPAR motifs ^29^ using the CAGEd-oPOSSUM web tool (Arenillas *et al*., in preparation). Merged sequences from 500 bps up‐ and down-stream of escTSSs were compared to two background sets: nonDT and a randomly selected %GC and length matched background set (escTSSs_bg). The TF binding motifs associated with YY1, FOXC1, ETV1, Foxo1 and Atf3 were found to be over-represented around escTSSs when compared to both background sets (Fig 3; S3 Table). The significance of YY1 binding motif was highest among all motifs, and it was 2.4 and 1.5 times more frequently found in escTSSs than escTSSs_bg and nonDT sets, respectively. In comparison, the other 4 motifs ranged between 1.4-1.6 and 1.2 fold higher than the background sets. The motif over-representation analyses indicated YY1 as a potential regulator of escapees through sequence specific binding.

**Figure 3:**
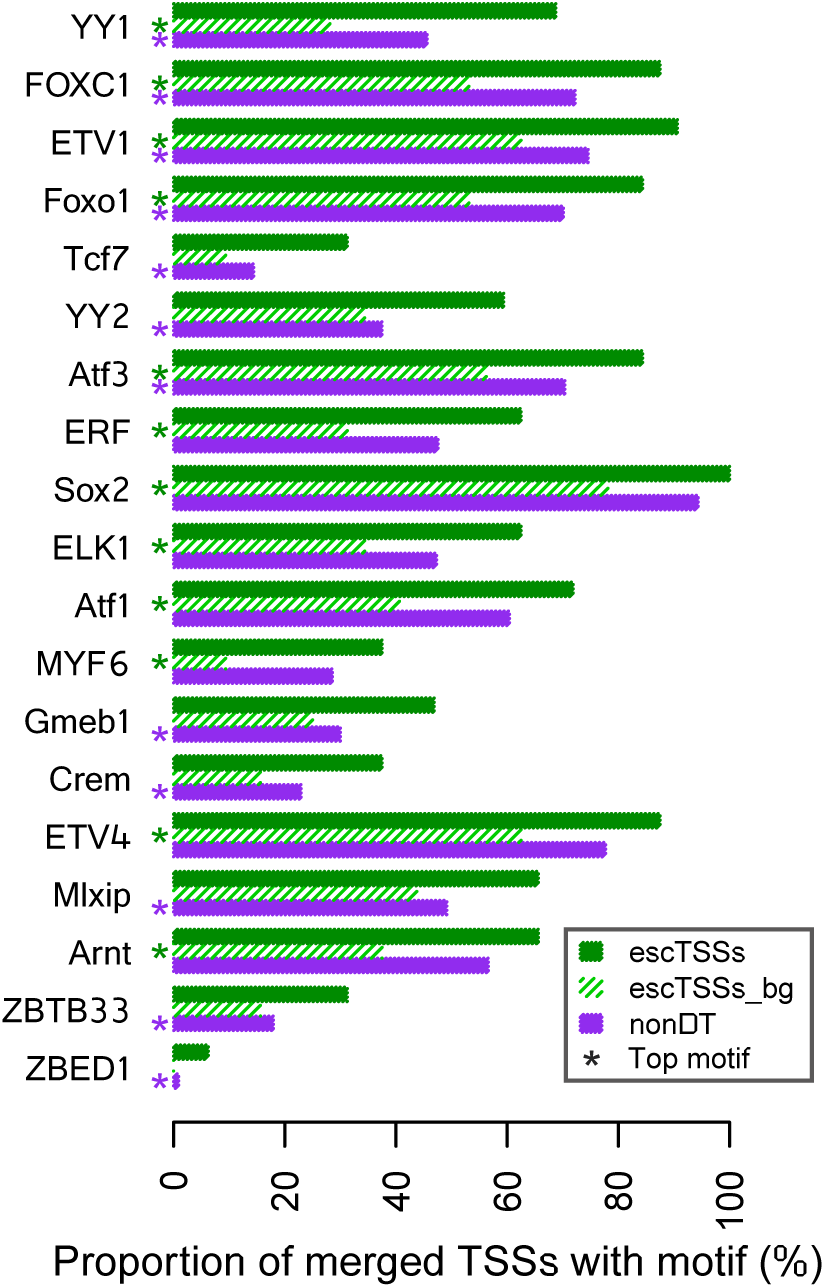
TF binding motif over-representation analyses of escape TSSs compared to two background sets accounting for different properties. Barplots showing the proportions of merged TSS regions (x-axis) containing the JASPAR TF motifs labeled on the y-axis for escTSSs (green) and non-differentially transcribed set on X with Bonferroni-corrected p-values equal to 1 (nonDT; purple). The proportions for %GC composition and length matched background set sampled from the genome for escTSSs (escTSSs_bg) is shown in shaded green. The top motifs with Fisher scores greater than 95th percentile in escTSSs compared to either background set (escTSSs_bg and nonDT) are plotted. The top motifs derived with each background set are marked with asterisks under the background bars with corresponding color. Motifs are presented in decreasing order of the sum of Fisher scores from both comparisons.

### Over-representation of TF ChIP-seq peaks around bi-allelically transcribed escTSSs

To explore the regulation of escapees at the experimental level, we examined TF ChIP-seq data in male and female cell lines from the ENCODE project to identify over-represented TF binding at the bi-escTSSs. TF binding on chrX in female cells can be either bi‐ or mono-allelic, and the measured binding degree would reflect a mixture of both X chromosomes. Therefore, given a positive-regulating TF, we expect a stronger binding pattern at bi-escTSSs than mono-allelically active TSSs on chrX in female cells (S1 Fig C). We refer to this expectation as the bi-allelic effect. As there is only one copy of the chrX in the male cells, enrichment is not expected.

We tested the over-representation of all 689 uniformly processed ENCODE TF ChIP-seq peak sets within 500 bps of bi-escTSSs compared to background TSSs on chrX (X_bg) and autosomes (Auto_bg) with matched average expression (see Methods for details). Twenty-seven individual ChIP-seq datasets, corresponding to 19 unique TFs, were overrepresented around bi-escTSSs when compared to X_bg (One-sided Fisher’s exact test, Bonferroni-corrected p-values ≤ 0.05; S4 Table; Fig 4A). The top 10 unique over-represented TFs in the order of decreasing significance were c-Myc, Pol2, TBLR1, PAX5-C20, YY1, CTCF, FOXM1, ATF2, SIN3A and CHD1. Notably, the significant TF datasets were all generated from female cells. Although non-significant after Bonferroni correction, YY1 was the highest TF ranked in multiple male cells. By contrast, when compared to the Auto_bg, none of the TFs were significantly over-represented (S4 Table and S3 Fig A). Taken together with the result from the motif over-representation section, the significance of YY1 binding in cellular context further highlighted the strong potential for a role for YY1 in facilitating the escape from XCI.

**Figure 4:**
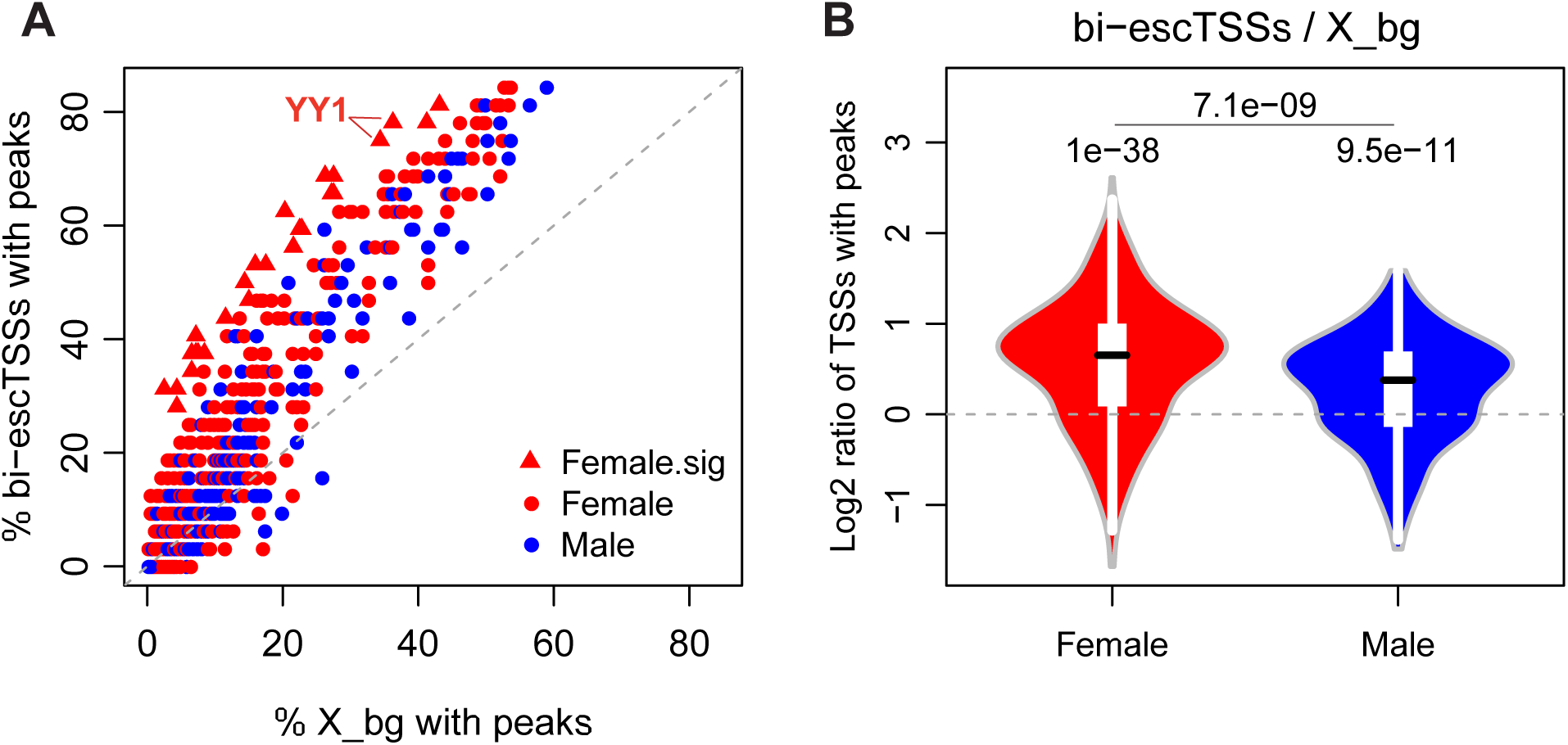
Over-representation of ENCODE TF ChIP-seq peaks at bi-escTSSs and differences between the sexes. (**A**) Scatter plot showing the percentages of TSSs that overlap a peak comparing between bi-escTSSs (y-axis) and matched background TSSs on chrX (X_bg; x-axis). Each dot represents a uniformly processed TF ChIP-seq dataset from ENCODE. Red and blue colors represent female and male cells in all plots, respectively. The dashed gray lines are the baseline reflecting no differences between proportions of escTSSs and X_bg overlapping peaks. Datasets with significant overrepresentation of peaks in escTSSs compared to X_bg are displayed as triangles (Bonferroni-corrected p-values ≤ 0.05). Significantly over-represented YY1 datasets are labeled on the figure. (**B**) Violin plots showing the distributions of log2 ratio of escTSSs to X_bg in female and male cells. The p-value from comparing the log2 ratios between male and female cells (one-sided Wilcoxon tests) and the p-values of one-sample Wilcoxon tests for the distributions are shown above the violin plots.

While the 27 ChIP-seq datasets have peaks significantly over-represented at escTSSs compared to X_bg, many additional sets (204 of 689) were reported non-significant after the conservative Bonferroni correction was applied. We observed an overall trend of higher proportions of escTSSs containing peaks compared to that of X_bg (Fig 4A). Indeed, the distribution of log2 ratios of percent overlaps with TF peaks (escTSSs/X_bg) in both female and male cells were significantly greater than 0 (one-sample Wilcoxon test p= 1.0*10^−38^ and 9.5*10^−11^, respectively; Fig 4B). In agreement with our expectation, the log2 ratios in female cells were significantly higher than that observed in male cells (onesided Wilcoxon test p= 7.1*10^−09^). The log2 ratios comparing escTSSs to autosomal background (Auto_bg) was not significantly different from 0 in female cells (p=0.51; S3 Fig B), whereas it was significantly lower in male cells (p=1.7*10^−18^), due to the single copy of chrX in male cells. The overall higher number of binding events at escTSSs stressed the importance of taking multiple TF datasets into consideration when comparing between sexes at escTSSs or between escTSSs and other chrX TSSs, otherwise the differential occupancy may merely reflect the bi-allelic effect.

### ChIP-seq read depth reveals overall reduction of input from heterochromatic Xi

As TF binding peaks do not necessary reflect linearly the magnitude of TF occupancy, we further compared the read depth at escTSSs to the same background sets using input and YY1 ChIP-seq data from two female and male cells. Although female cells have two copies of autosomes and chrX, the read depth for input within ± 50 bps of X_bg was 0.61 fold lower than that of Auto_bg on average, while the read depth within ± 50 bps of escTSSs was closer to Auto_bg (0.95 fold; Fig 5A). This indicated a discounted number of input fragments available for capture due to the compactness of Xi around X_bg in female cells. While male cells have two copies of each autosome and one of chrX, the input read depths at chrX (X_bg and escTSSs) and autosomes (Auto_bg) reflected the ratio as expected (0.48 and 0.46 fold, respectively; Fig 5B). Overall, the input reads in female cells reflect bi- and mono-allelic activities at escTSSs and X_bg, respectively, and despite the mono-allelic activities at X_bg in female cells, input reads at X_bg in female cells were higher than those in male cells, reflecting a reduction of the heterochromatin input.

**Figure 5:**
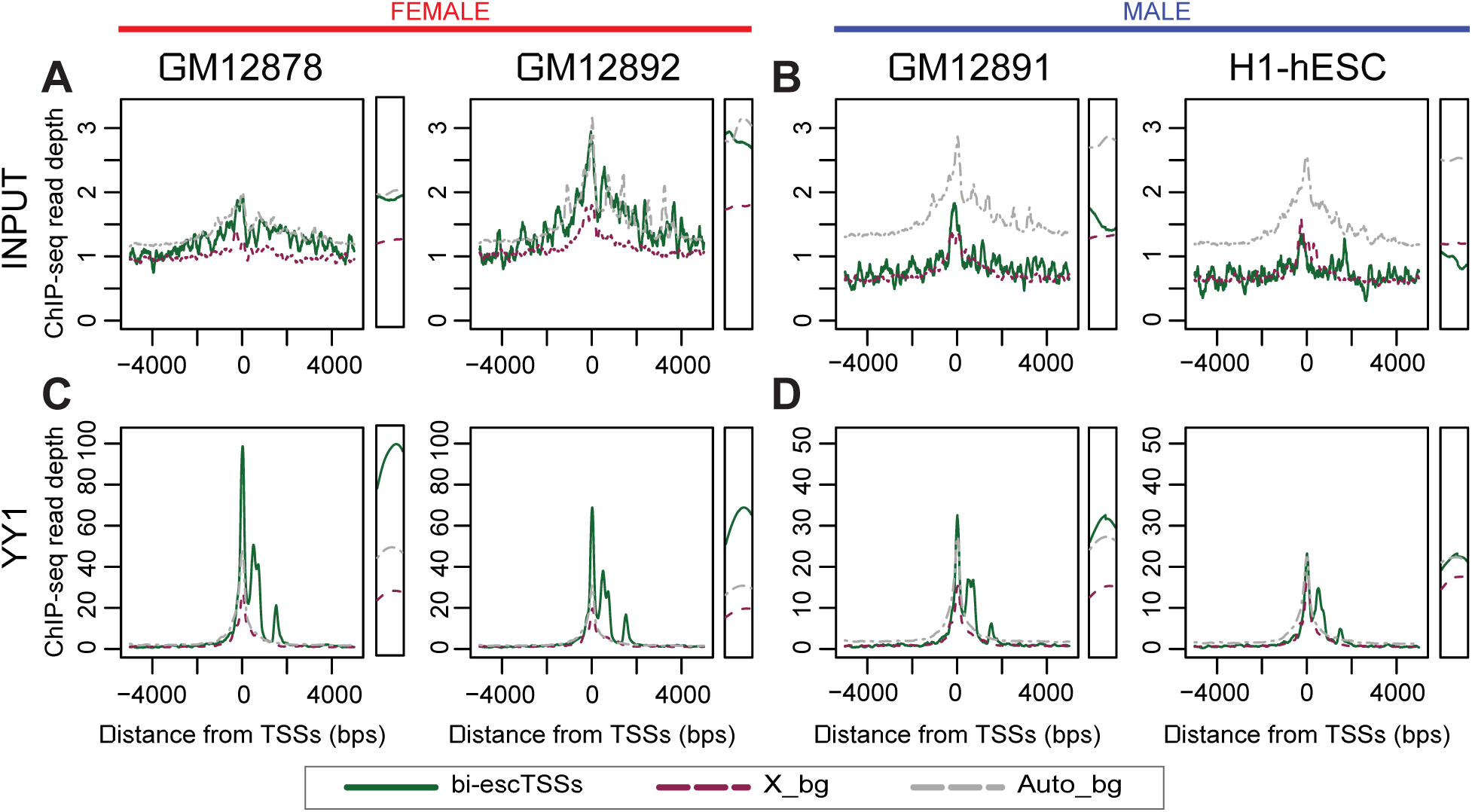
Input and YY1 ChIP-seq read depths around bi-escTSSs for both sexes. The read depth plots for ENCODE ChIP-seq input samples in two female (**A**) and male cell lines (**B**) within 5kb of three TSS sets: a subset of bi-escTSSs in green with a filter of unique escTSS per gene symbol, background TSSs on chrX with matched averaged expression (X_bg) in violet dashed line, and autosomal TSSs with matched averaged expression (Auto_bg) in gray dashed line. The read depth plots for ENCODE YY1 ChIP-seq data in the same female (**C**) and male cells (**D**). The scales on y-axis of figure A and B are the same, while the scale of C is two times the scale of D to reflect the expected X-copies in female and male cells. For the ease of visualization, the narrow panel on the right of each plot displays the read depth within 50 bps of the TSSs.

### YY1 binding at escTSSs shown by read depth and allelic analyses

Interestingly, despite slightly lower input levels, YY1 ChIP-seq read depth at escTSSs was 2.1 times that of Auto_bg in female cells (Fig 5C). Furthermore, despite the 1 to 2 ratio of input levels at escTSSs and Auto_bg in male cells, the YY1 read depths at escTSSs and Auto_bg were similar (1.1 fold; Fig 5D). In contrast, the YY1 read depth ratio between X_bg and Auto_bg was similar to that of input read depth in both female and male cells (Figs 5C and 5D). Overall the results showed higher occupancy of YY1 at escTSSs compared to both X_bg and Auto_bg for both sexes, and thus enhanced YY1 binding may predispose to ongoing transcription from the inactive X.

Given that YY1 motifs and ChIP-seq peaks were over-represented at escTSSs, we next examined the allelic binding of YY1 on chrX in the female GM12878 cell line to probe its functional role, and confirm its bi-allelic binding at bi-escTSSs (see also schematic in S1 Fig D). We extracted the YY1 ChIP-seq read counts on Xi and Xa at the 67 heterozygous sites within YY1 ChIP-seq peaks in GM12878 (S5 Table). Only two YY1 peaks containing heterozygous sites were within 500bps to two bi-escTSSs. Indeed, YY1 was bound on both Xa and Xi at both sites, within exon 1 of *SMC1A* (7:13, representing counts of Xa:Xi) and exon 1 or exon 2 of *JPX* transcripts (12:8), confirming bi-allelic transcription and bi-allelic YY1 binding of the escapees.

Consistent with previous reports of Xa-biased binding of TFs reflecting Xa-biased transcription in female cells, the majority of heterozygous sites (52 out of 67) had more reads on Xa than Xi, while 13 heterozygous sites had more reads on Xi. Despite the overall trend of Xa-biased binding, we found the highest number of allelic YY1 reads on Xi (3:474) within exon 1 of *XIST*. The second highest number of YY1 reads was at 215 bps upstream of *UPF3B* (140:0), and the Xa-biased binding agreed with its subject status reported previously ^11^. The site with the third highest number of YY1 reads overlapped intron 5 of *FIRRE* and was Xi-biased (4:138). Interestingly, Xa-biased YY1 was detected at sites 477 bps upstream of *FIRRE* TSS and in intron 1 (44:5 and 20:0). Such discrepancy was consistent with the previous report of a shorter alternative *FIRRE* transcript on Xi ^4^ and the finding of female-specific enhancers in introns 2-12 from ATAC-seq^12^. As YY1 can be bound specifically on either Xa or Xi, there likely are cofactors acting at escapees in collaboration with YY1. Overall, the allelic binding reads of YY1 indicated a positive association with regulation of X-linked genes.

### Significant Xi-biased YY1 occupancy at *XIST*, *FIRRE* and two other superloop-associated lncRNAs in GM12878

Given the important roles of *XIST* and *FIRRE* on XCI and the strong allelic imbalance of YY1 binding at these lncRNAs, log2 ratios of Xa to Xi counts were computed to identify strong allelic imbalance at all 67 heterozygous sites (see Methods for detail). The significant correlation between allelic imbalance scores from replicates of YY1 ChIP-seq datasets indicated strong consistency (ρ=0.87 with p < 2.2*10^−16^; Fig 6). Heterozygous sites at exon 1 of *XIST* and intron 5 of *FIRRE* were shown to have the strongest Xi-biased YY1 binding. These sites with YY1 binding were found to be top 5 in significance of allelic imbalance when we extended the allelic analysis to all available ChIP-seq and DNase I datasets in GM12878 as well as 1,321 heterozygous sites, and tested each heterozygous site and data pairs for imbalance significance (see Methods for details; raw reads in S6 Table). We identified 389 significant imbalance site-data pairs comparing to the overall Xa-biased norm using False Discovery Rate (FDR) correction (discussed in S1 File; S4 Fig), which corresponded to 178 unique heterozygous sites (FDR corrected p-values ≤ 0.05; S7 Table). Thirty-eight out of 178 sites exhibited imbalance in more than one dataset, within which 16 and 18 sites were biased consistently across data sets towards Xi and Xa, respectively. Such consistency in allelic imbalance across TFs indicated broader influence such as open or closed chromatin rather than individual TF binding affinity of alleles.

**Figure 6:**
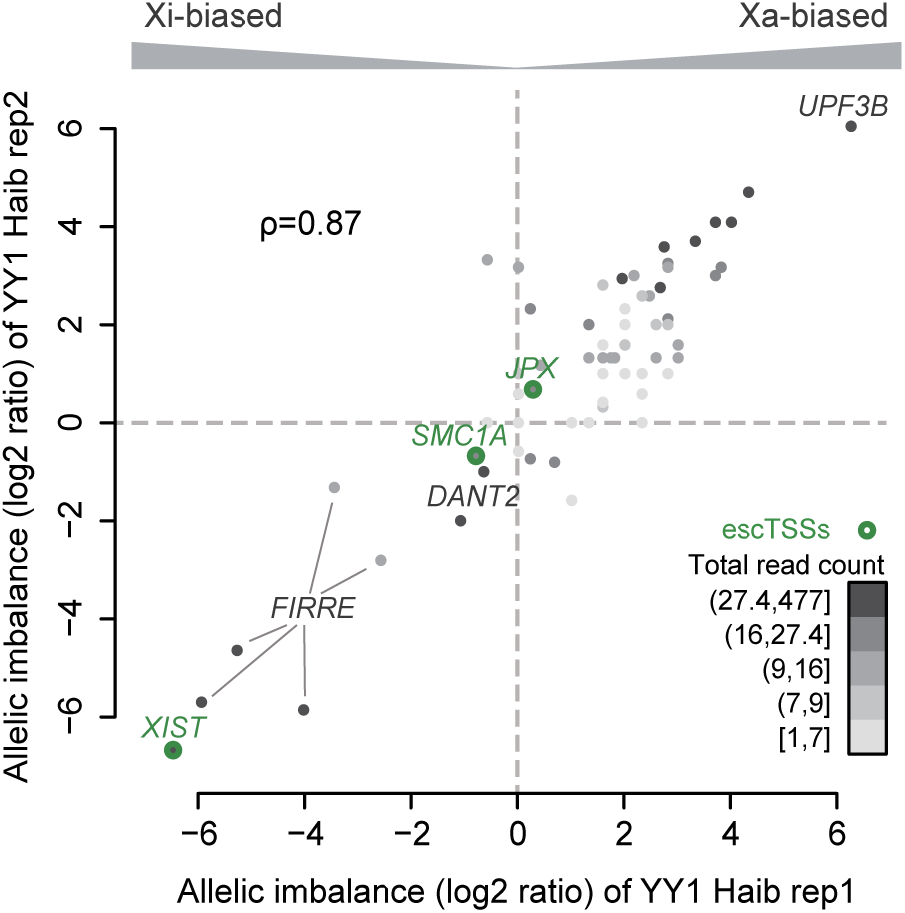
Allelic imbalance at heterozygous sites within YY1 ChIP-seq peaks on chrX in the GM12878 cell line for two replicate samples. Scatterplot showing the allelic imbalance of replicated YY1 ChIP-seq data sets from ENCODE (see Methods). For visualization purposes, allelic imbalance is represented by the log2 ratio of (Xa+1) to (Xi+1). A positive log2 ratio value indicates more reads on Xa, while a negative value represents more reads on Xi. Each of the 67 dots represents a heterozygous site within a YY1 binding peak. The Pearson correlation between allelic imbalance of the replicated datasets is 0.87. Dotted lines indicate the baselines for balanced allelic binding. Heterozygous sites within 50 bps of escTSSs are indicated by green circles. The intensity of shading of each dot reflects the total number of YY1 reads from both replicates at the heterozygous site (where read counts were assigned to five 20 percentile bins).

Interestingly, the majority of significantly Xi-biased imbalances mapped to the vicinity of the four lncRNAs associated with Xi-specific superloops (Fig 7; S7 Table). YY1 binding at the heterozygous site within *XIST* was the most significantly Xi-biased across all data sets and sites (FDR-corrected p-value= 7.12*10^−221^), while 21 other datasets were also significantly Xi-biased at the same site to a lesser degree. The Xi-biased YY1 binding at heterozygous sites around intron 5 of *FIRRE* were ranked fourth and fifth in overall significance, whereas RAD21 and CTCF were preferentially Xi-bound with lower ranks. While no heterozygous sites overlapped *DXZ4*, significant Xi-biased sites overlapped the *DXZ4* associated non-coding transcript 2, *DANT2*. Notably, *DANT2* is located within the previously reported boundary of two Xi superdomains, which were not associated with Xa or in male cells ^8^. Overall, significant Xi-biased bindings of YY1 and MYC were found in all four lncRNAs, while RAD21 and CTCF were significantly Xi-biased at *FIRRE* and *LOC550643*, but not *XIST* and *DANT2*. The allelic analysis revealed Xi binding of YY1 at Xi-transcribed genes in general, including escTSSs and key lncRNAs involved with XCI, Xi-specific superloops and the superdomain boundary.

**Figure 7:**
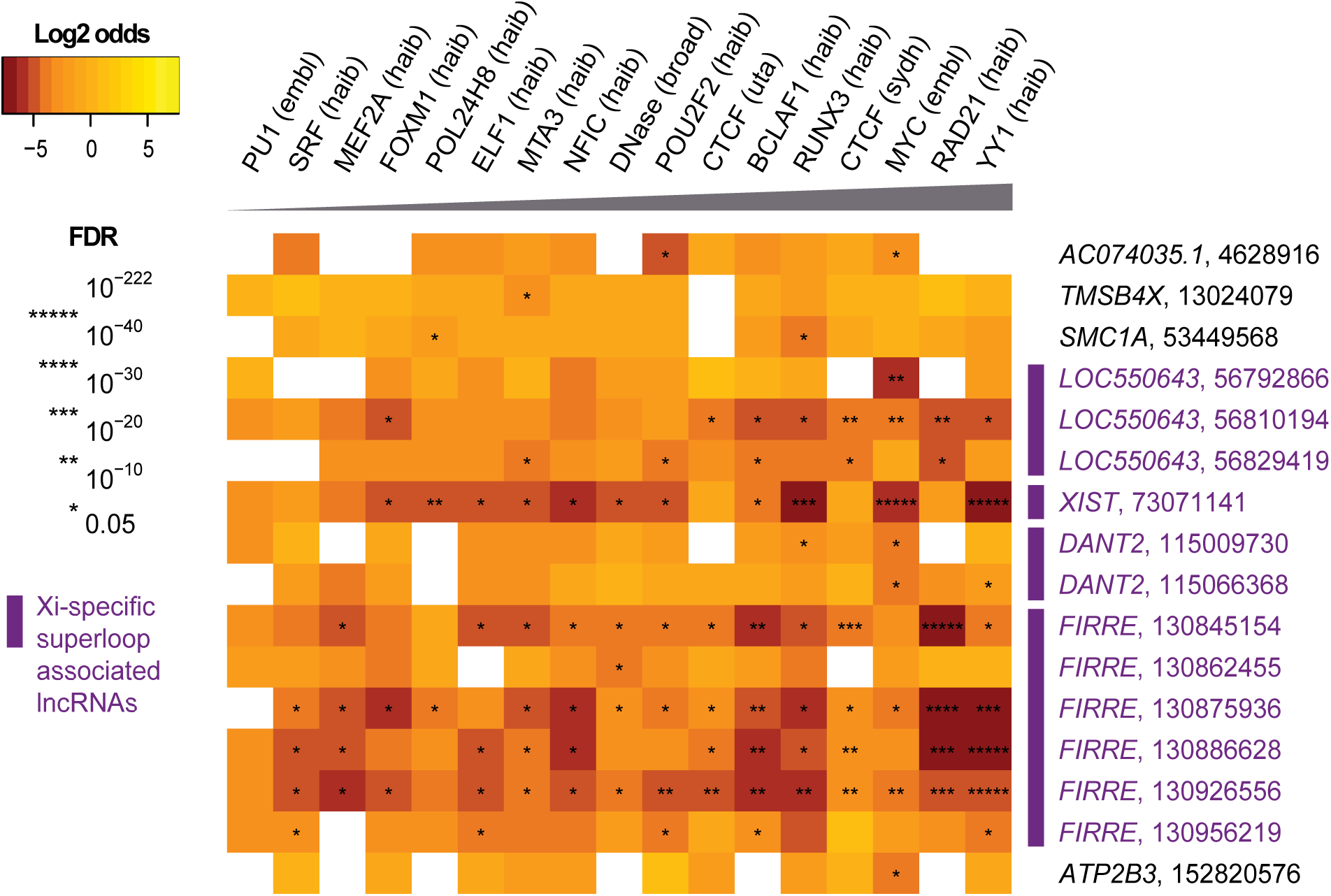
Heterozygous sites with significant allelic imbalance towards Xi observed in multiple datasets from the GM12878 lymphoblastoid cell line. Heatmap of heterozygous sites (rows) significantly Xi-biased in more than one dataset (ChIP-seq and DNase I data; column). Only datasets that are significantly Xi-biased at more than four heterozygous sites are listed. Datasets are denoted by the feature name followed by the ENCODE lab where data was generated, and heterozygous sites are denoted by the gene name of the nearest TSS followed by the chrX coordinate of the site. Colors in the heatmap represent log2 odds ratio values reflecting Xa‐ or Xi-biased binding of the TF with positive (gold) or negative (brown) values, respectively. The log2 odds ratio distinguishes Xi bias (negative) and Xa bias (positive). White boxes indicate zero read counts at the corresponding site-data pair. The degrees of significance estimated by FDR-corrected p-values are indicated on a scale of 1 to 5 asterisks with the corresponding p-value thresholds shown in the legend. The datasets from left to right are ordered in increasing counts of higher significance scales denoted by the gray triangle, and the heterozygous sites are ordered using genomic coordinates on chrX. The four lncRNAs previously reported to be associated with Xi-specific superloops are marked with purple bars and colored in purple.

## Discussion

Here we analyzed CAGE datasets reporting the transcription levels at chrX TSSs to identify escTSSs through differential transcription analysis between male and female samples matched for cell categories. Predicted escTSSs yielded the highest precision and the second highest recall for escapee gene identification compared to three reports using different techniques. Analyses of experimental data revealed the unique properties of escapees and the resulting bias of bi-allelic activity in data, and identified the potential involvement of YY1 binding. Allelic binding in GM12878 cells further highlighted a role for YY1 not only at escTSSs but also proximal to lncRNAs involved with Xi-specific superloops. In aggregate, the analyses indicated that YY1 influences Xi expression of not only XIST but more broadly of escapees.

From a bioinformatics perspective, the analysis of chrX is challenging. In female datasets, measurements captured are a combination of properties from Xa and Xi. Genes subject to XCI are under a mixture of positive (Xa) and negative (Xi) regulation, whereas bi-allelically transcribed escapees are positively regulated on both X chromosomes. Direct use of tools and approaches designed for autosomes can confound interpretation. Our analyses explored properties of bi-escTSSs both in DNAm and TF binding perspectives. Despite having higher expression in females, bi-escTSSs have similar total DNAm levels between sexes consistent with bi-allelic activity in females. In ChIP-seq datasets, there are more bi-escTSSs with peaks than the mono-allelically active background TSSs on chrX for both sexes (recall Fig 4B). Such over-representation in female datasets resulted from a combination of the bi-allelic effect, and predispositions of TFs to bind at escapees. Taking all data into consideration, the ChIP-seq peaks of 19 TFs, including MYC and YY1, which were significantly over-represented at bi-escTSSs in female cells indicated functional roles at escapees with potential predisposition beyond the bi-allelic effect. We have revealed properties of escapees by addressing chrX with targeted analyses of genome-scale data.

The ubiquitous TF YY1 is a transcription initiator with diverse cellular functions ^30^. Motif and ChIP-seq peak enrichment at bi-escTSSs and allelic binding highlighted the role of YY1 binding in positive regulation of genes on Xi. Connections between YY1 and *XIST* expression have been previously reported. A study that knocked down 59 TFs and chromatin modifiers in a HapMap lymphoblastoid cell line ranked YY1 second, after IRF4, in significantly affecting the expression of *XIST*^31^. Furthermore, YY1 was reported to support transcription of *XIST* through tethering RNA ^6,7^, and its capability to bind both DNA (on Xi) and RNA of *XIST* made it crucial for *XIST* localization at the nucleation center and maintenance of XCI ^5^. A recent report on enhanced YY1 occupancy through trapping by RNA in mouse also provided relevant insights on the mechanism, which leads to stability in gene expression ^32^. Given the over-representation of YY1 motif and peaks as well as its Xi binding at escapees, our result suggests that YY1 plays a broader role in the regulation of escapees. Its trapping by RNA provides a possible mechanism for facilitating escape from XCI, which can be further experimentally examined through interactions between YY1 and escapee RNAs.

YY1 has been implicated in controlling long distance DNA interactions, and controlling non-coding antisense RNA transcripts in B cell development ^33,34^ Taken together with the significant YY1 Xi-biased binding at the four lncRNA loci where Xi-specific superloops are highly interactive, the evidence supports a potential role of YY1 for mediating long distance DNA interactions involving Xi. While CTCF and RAD21 have been prominently studied as mediators of chromatin looping ^8^, there may be interplay between YY1 and CTCF for Xi transcription, as peaks of both TFs are enriched at boundaries of topologically associating domains and nuclear compartments ^35^. Furthermore, YY1 and CTCF co-bound regions are transcriptionally active and highly conserved in primate genomes, and have been suggested to act together within the chromatin remodeling machinery for activation of euchromatin regions ^36^. A lack of CTCF motif and peak over-representations at escTSSs as well as the absence of CTCF-binding tandem repeats at *XIST* ^8^ suggest a broader role of YY1 in looping at both bi-allelic and mono-allelic escapees. As active regions on Xi loop apart from the condensed Xi ^37-39^, YY1 may be functionally contributing to the phenomena.

Since YY1 is a ubiquitous TF and exhibited Xa-biased binding at genes subject to XCI, it is likely working with other TFs in regulating escapees. We also identified motif and peak over-representation of forkhead (FOX) transcription factors at escTSSs, but did not find evidence of interaction with YY1 in the literature. Over-representation of MYC ChIP-seq peaks in proximity to escTSSs as well as allelic imbalance at the superloop-associated lncRNAs also suggested a potential role of MYC in facilitating escape from XCI. MYC is a key regulator of cell proliferation involved with the release of promoter-proximal pausing of RNAPII ^40,41^. YY1 has been reported to be an active component of the MYC transcription network in mouse embryonic stem cells ^42^. Since the MYC binding motif was not over-represented at escTSSs, the results suggest that YY1 works collaboratively with MYC in regulating escapees through YY1 sequence-specific binding. Interestingly, both *YY1* and *MYC* have been reported to be overexpressed in numerous cancers ^43,44^, and a recent study highlighted the epigenetic instability of the inactive X in breast cancer cells leading to X-linked gene reactivation ^45^. Further studies will be required to examine the connection between these TFs and cancer in the context of the X chromosome, and potentially elucidate sex differences in cancer susceptibility and progression.

Overall the results support a model in which YY1 facilitates escape from XCI, but the analyses are generally limited by the availability of data. At transcription activity level, our approach in identifying differentially transcribed TSSs from FANTOM5 CAGE datasets was limited to ubiquitous escapees, as there are few matched male and female samples. Further investigation of relationships between cell categories and sexes could enable the identification of cell type-specific escapees, but will require sufficient numbers of samples from both sexes in the same cell type. At TF binding level, both the JASPAR motif collection and the ENCODE ChIP-seq datasets are far from complete, covering only a subset of TFs. Furthermore, the allelic imbalance analysis in human is particularly constrained to a small set of heterozygous sites within TF peaks and within the female cell lines that are skewed in XCI. Lastly, our allelic analysis on chrX did not take into account the TF binding affinities towards different alleles, as we intended to capture the epigenetic influence from XCI, which was supported by the observation of multiple TFs sharing significant Xi-biased binding at individual sites. Despite the limitations, we were able to reveal the intrinsic data properties of X-linked genes in males and females, which in turn allowed the identification of the key regulator of escapees and its functional role on the inactive X.

## Methods

All analyses were conducted in R (3.0.2) ^46^ and Bioconductor (2.13) ^47^ unless otherwise stated.

### Public datasets

For expression analysis and sex classification, the RLE-normalized expression table of robust CAGE peaks for human samples was retrieved from the FANTOM5 consortium ^17^For the DNA methylation analysis, raw data files in idat format from Illumina 450k array generated by TCGA were retrieved ^28^, and the following datasets for four cancer types were used: Bladder Urothelial Cancer (BLCA), Colon Adenocarcinoma (COAD), Head and Neck Squamous Cell Carcinoma (HNSC) and Lung Adenocarcinoma (LUAD). For the ChIP-seq and DNase I analysis, uniformly processed ChIP-seq peaks and DNase I peaks were retrieved from the ENCODE ^20^ page on UCSC genome browser. For generating read depth plots, normalized bigwig files from ENCODE were retrieved from https://sites.google.com/site/anshulkundaie/proiects/wiggler. For allelic binding analysis, a total of 213 files of ChIP-seq, input and DNase I data from GM12878 cells were retrieved from the ENCODE ^20^ page on UCSC genome browser.

### Classification and differential expression of the sexes using FANTOM5 CAGE data

All CAGE TSSs were labeled with the gene name corresponding to the nearest TSSs of Ensembl transcripts from ENSEMBL GENES 71 ^48^. A supervised Random Forest classifier using randomForest R package ^49^ was implemented to classify the sexes for the FANTOM5 CAGE data sets. Data from samples involving treatments of the cells or tissues were excluded, resulting in a total of 809 datasets. Of these, 296 male and 234 female labeled samples were used as training data. Two classifiers were constructed, one using 5071 TSSs in the non-PAR portions of chrX as features, and the other using the 56 *XIST*-associated TSSs. Performance of the classifier was assessed using 10-fold cross validation as well as out of bag error estimates. For the selected 5,071 TSS-based classifier, sample outliers were identified based on out-of-bag error estimation. The classifier was used to predict samples labeled unknown.

Ontologies associated with the FANTOM5 samples were retrieved from FANTOM5, including Cell Ontology, Uberon, and Disease Ontology. One hundred and fifty-three cell category terms with association to at least 30 samples were extracted. To take cell categories into account when assessing differential transcription between sexes, the first 29 principal components of cell categories explaining over 90% variance in addition to sexes were used as covariates of the linear regression model: 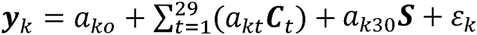, where ***y***_*k*_ is the vector of log10 (tags per million counts at TSS *k* +5) values in all samples reflecting the level of transcription, ***C***_*t*_ is the vector of rotated cell category relevance values of each sample for the *t*^th^ principal component, and ***S*** is the vector of sexes of samples from the sex classifier. The fitted coefficient of the sex variable, *a*_*k*30_, and its p-value were used to assess differential transcription between sexes of TSS *k*. TSSs with Bonferroni-corrected p-values less or equal to 0.05, and negative *a*_*k*30_ values (higher expression in female samples) were defined as escTSSs. The precision and recall values compared to escapee lists from the literature were computed using the count of true predicted escapees over the count of all predicted escapees, and the true escapee count, respectively. Genes reported in more than one list were taken to constitute true escapees. Fig 1C is generated using the VennDiagram R package ^50^.

### Similarity and differential analysis of DNA methylation between sexes

Due to the complication of XCI in female samples, a basic preprocessing approach using background subtraction of negative control probes and internal control normalization was conducted on all Illumina 450k DNA methylation data from TCGA using the Minfi R package ^51^. The beta value, *β*_*i*_, representing the degree of DNAm was computed for each probe *i* as *Meth*_*i*_/(*Meth*_*i*_ + *Unmeth*_*i*_ + 100) by default. We adopted the analysis of microarray studies by computing the logged differential methylation value between sexes (*M*) and the logged average methylation value (*A*) for each probe 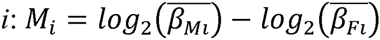 and 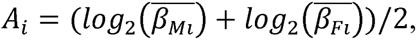, where 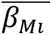 and 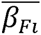 were the average *β* values of probe *i* for the male and female samples, respectively. For probes on chrX, we fitted a linear model,***M*** = *d*_0_ + *d*_1_***A***,byrobust regression using the estimating function of Tukey’s bisquare, which gave extreme observations zero weights. We then obtained the residual of each probe to be the DNAm similarity score between sexes. For assessing the overall correlation between DNAm similarity scores and differential transcription of CAGE TSSs, we assigned probes within 50bps to each TSS. When multiple DNAm probes are within 50bps of a TSS, the average of the similarity scores was computed. For probes on autosomes, differential methylation analysis was performed, including age and cancer stage as covariates (adapted from the dmpFinder method of Minfi). F statistics were computed by comparing the goodness of fit of the models using age and cancer stage as covariates with and without the addition of sex variable. Bonferroni-corrected p-values were obtained for each autosomal probe.

### Motif over-representation tests of escTSSs using CAGEd-oPOSSUM

The escTSSs were subjected to motif over-representation analysis using our CAGEd-oPOSSUM web tool (Arenillas *et al*., in preparation: http://cagedop.cmmt.ubc.ca/CAGEd_oPOSSUM/). A total of 478 JASPAR2016 motifs with a minimum specificity of 8 bits for vertebrates ^52^ were used for the prediction of binding sites. The “Use only FANTOM5 CAGE peaks identified as true TSSs by the TSS classifier” option was selected to filter out CAGE peaks that were less likely to be TSSs. The remaining CAGE peaks were extended 500 bps up‐ and down-stream, and merged. Two tests were conducted differing in background sets: (i) random CAGE TSSs with %GC composition and length matched to escTSSs sampled by CAGEd-oPOSSUM, and (ii) 4,939 non-differentially transcribed chrX TSSs with Bonferroni-corrected p-values of 1 from the differential transcription analysis. The TF binding score threshold was set to 85% by default, and the Fisher scores for both tests were retrieved to assess the over-representation significance of each motif around escTSSs.

### TF ChIP-seq peak over-representation testing and read depth plots

Over-representation testing of TF ChIP-seq peaks within 500bps of bi-escTSSs was conducted using the one-sided Fisher’s exact test. To avoid double counting, only one TSS with the strongest differential expression between sexes was selected per escape gene. As a result, we conducted the analysis on 32 bi-escTSSs, and randomly selected 320 background TSSs from chromosome X (X_bg) and 3200 from autosomes (Auto_bg) with matched average transcription levels. An additional criterion of Bonferroni p-value equal to 1 (from differential transcription analysis) was used for the X_bg set to avoid selecting borderline escapees. We assigned all CAGE TSSs to 5 equally sized bins according to average expression percentile ranks. The percentage of bi-escTSSs assigned to each bin was determined and a background set was selected to match the distribution across bins. We extracted the counts of bi-escTSSs, X_bg and Auto_bg sets having at least one peak within 500bps for each ChIP-seq data. Given the overlapping and non-overlapping counts of bi-escTSSs and background TSSs, one-sided Fisher’s exact tests were conducted to test for positive association between the peaks of each ChIP-seq data and the bi-escTSSs. The significance was adjusted for multiple testing using Bonferroni correction. The log2 ratios of bi-escTSSs to each background set with peaks were computed as log2(% bi-escTSSs with peaks/ % background with peaks). For the read depth plots, Bwtool ^53^ was used to extract the YY1 ChIP-seq and input read depth within 5kb of the three TSS sets from big wig files (bi-escTSSs, X_bg and Auto_bg). Read depth ratio between any two TSS sets were computed by comparing the average read depths within ±50 bps of the TSSs. The ratios from cells of the same sex were averaged.

### Allelic ChIP-seq and DNase I data analysis of the female GM12878 cell line

Our in-house allelic binding pipeline was used to extract reads at heterozygous sites from GM12878 datasets and assess mapability for filtering (Shi *et al*., in preparation). We obtained the genotype data of GM12878 from the 1000 Genomes Project ^54^, and a personalized hg19 genome for GM12878 was built by representing single nucleotide variations as degenerate IUPAC codes. Raw reads of 214 ChIP-seq, input and DNase I datasets were aligned to this personalized genome, using Novoalign (version 3.01.00: http://www.novocraft.com) with default parameters. The phased information of GM12878 was obtained from Illumina (http://www.illumina.com/platinumgenomes/). As allelic mapping bias may exist at certain locations within the personalized genome, a set of simulated reads were generated for mapability assessment. We first merged all GM12878 TF ChIP-seq binding peaks plus 100 bps flanking regions, and then simulated equal numbers of reads for each allele of the single nucleotide variants to map to the personalized genome. Only 1,321 out of 1,497 heterozygous sites with balanced simulated read counts between two alleles, calculated by requiring the read count of either allele divided by the sum of two alleles to be between 0.6 and 0.4, were kept for further analysis. As paternal X chromosome corresponds to the Xi in GM12878, we assigned the allelic read counts to Xi or Xa at each heterozygous site accordingly.

All replicated data sets generated by the same ENCODE lab were merged through summing Xi and Xa reads separately at each site, resulting in 101 merged data sets. The only exception was for Fig 6, where replicated YY1 datasets generated by the HudsonAlpha Institute for Biotechnology lab (Haib) were displayed separately to show the correlation of allelic imbalance between replicates. For the individual allelic YY1 binding analysis, only 67 heterozygous sites within uniformly processed YY1 ChIP-seq peaks were examined to avoid noise from low occupancy. Allelic imbalance score at site *h* was computed for visualization purposes using the log2 ratio of reads at Xa over Xi with a constant of 1 added to avoid zero denominators: *log*_2_((*Ra*_*h*_ + 1)/(*Ri*_*h*_ + 1)), where *Ra*_*h*_ and *Ri*_*h*_ represented read counts on Xa and Xi at site *h*, respectively.

For the batch analysis of allelic imbalance, all 1321 sites were used. The significance of allelic imbalance was assessed using Fisher’s exact test comparing the Xa and Xi counts at each heterozygous site to the total Xa and Xi counts within the corresponding data. For each data *g* at site *h*, the 4 values in 2x2 table were 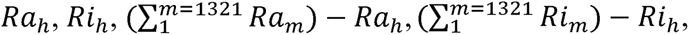 where *Ra*_*h*_ and *Ri*_*h*_ represented read counts on Xa and Xi at site *h* from data g, respectively. The significance reflected the extremity of imbalanced site-data pairs compared to the overall Xa-biased total read counts. With the 1321 sites and 101 merged data sets, a total of 61,470 site-data pairs with non-zero read counts from both Xi and Xa were tested. FDR p-value correction was used here because we expected multiple hits of the same TF to multiple heterozygous sites overlapping the same gene. The log2 odds ratio was computed to reflect the direction of imbalance bias with a positive or negative value reflecting bias towards Xa or Xi, respectively: *log*_2_((*Ra*_*h*_ + 1) * *1m=1321 Rim—Rih/(Rih+1)/(1m=1321Ram—Rah)*.

## Acknowledgements

We thank members of the Wasserman and Brown labs for helpful comments and discussion, Dora Pak for management support, and Miroslav Hatas for systems support, and. We would like to thank all members of the FANTOM5 consortium for contributing to generation of samples and analysis of the dataset and thank GeNAS for data production.

This research was supported by the Canadian Institutes of Health Research Operating Grant (MOP-119586; www.cihr-irsc.gc.ca) to the Wasserman and Brown labs; the Genome Canada Large Scaled Applied Research Grant (174CDE; www.genomecanada.ca) to the Wasserman lab; the Postgraduate Scholarships-Doctoral Program from Natural Sciences and Engineering Research of Canada (PGS D; www.nserc-crsng.gc.ca), and the Four Year Doctoral Fellowship from the University of British Columbia (www.ubc.ca) to CYC; a PhD fellowship from China Scholar Council (201206110038; www.csc.edu.cn) to WS; the Child and Family Research Institute, and the British Columbia Children's Hospital Foundation to AM.

FANTOM5 was made possible by a Research Grant for RIKEN OSC from MEXT (www.mext.go.jp) to Yoshihide Hayashizaki, Grant from MEXT for the RIKEN PMI YH, Grant of the Innovative Cell Biology by Innovative Technology (Cell Innovation Program) from the MEXT to YH and Grant from MEXT to the RIKEN CLST.

## Author contributions

CYC, WWW, and CJB designed and conceived experiments. YL (sex classification), AMM (DNAm and XCI), AM (motif and ChIP-seq over-representation), and WS (allelic binding) contributed intellectually within their areas of expertise. CYC conducted all analyses. WS provided the allelic and simulated binding reads. AM and DJA developed and provided support for the CAGEd-oPOSSUM web server. The FANTOM Consortium contributed in generation of samples and analysis of the FANTOM5 CAGE data sets: MI (data production); HK (data handling management); TL (tag mapping); PC, YH, and ARRF (FANTOM5 management and concept). CYC, WWW, and CJB wrote the manuscript. All authors read and approved the final manuscript.

## References

1. Clayton, J. A. & Collins, F. S. Policy: NIH to balance sex in cell and animal studies. Nature 509, 282–283 (2014).

2. Cerase, A., Pintacuda, G., Tattermusch, A. & Avner, P. Xist localization and function: new insights from multiple levels. Genome Biol 16, 166, doi: 10.1186/s13059-015-0733-y (2015).

3. Hacisuleyman, E. et al. Topological organization of multichromosomal regions by the long intergenic noncoding RNA Firre. Nat Struct Mol Biol 21, 198–206, doi: 10.1038/nsmb.2764 (2014).

4. Yang, F. et al. The lncRNA Firre anchors the inactive X chromosome to the nucleolus by binding CTCF and maintains H3K27me3 methylation. Genome Biol 16, 52, doi: 10.1186/s13059-015-0618-0 (2015).

5. Jeon, Y. & Lee, J. T. YY1 tethers Xist RNA to the inactive X nucleation center. Cell 146, 119–133, doi: 10.1016/j.cell.2011.06.026 (2011).

6. Chapman, A. G., Cotton, A. M., Kelsey, A. D. & Brown, C. J. Differentially methylated CpG island within human XIST mediates alternative P2 transcription and YY1 binding. BMC Genet 15, 89, doi: 10.1186/s12863-014-0089-4 (2014).

7. Makhlouf, M. et al. A prominent and conserved role for YY1 in Xist transcriptional activation. Nat Commun 5, 4878, doi: 10.1038/ncomms5878 (2014).

8. Rao, S. S. et al. A 3D Map of the Human Genome at Kilobase Resolution Reveals Principles of Chromatin Looping. Cell 159, 1665–1680, doi: 10.1016/j.cell.2014.11.021 (2014).

9. Human genomics. The Genotype-Tissue Expression (GTEx) pilot analysis: multitissue gene regulation in humans. Science 348, 648–660, doi: 10.1126/science.1262110 (2015).

10. Hall, E. et al. Sex differences in the genome-wide DNA methylation pattern and impact on gene expression, microRNA levels and insulin secretion in human pancreatic islets. Genome Biol 15, 522, doi: 10.1186/s13059-014-0522-z (2014).

11. Cotton, A. M. et al. Landscape of DNA methylation on the X chromosome reflects CpG density, functional chromatin state and X-chromosome inactivation. Hum Mol Genet 24, 1528–1539, doi: 10.1093/hmg/ddu564 (2015).

12. Qu, K. et al. Individuality and variation of personal regulomes in primary human T cells. Cell Syst 1, 51–61, doi: 10.1016/j.cels.2015.06.003 (2015).

13. Cotton, A. M. et al. Analysis of expressed SNPs identifies variable extents of expression from the human inactive X chromosome. Genome Biol 14, R122, doi: 10.1186/gb-2013-14-11-r122 (2013).

14. Ding, Z. et al. Quantitative genetics of CTCF binding reveal local sequence effects and different modes of X-chromosome association. PLoS Genet 10, e1004798, doi: 10.1371/journal.pgen.1004798 (2014).

15. Kucera, K. S. et al. Allele-specific distribution of RNA polymerase II on female X chromosomes. Hum Mol Genet 20, 3964–3973, doi: 10.1093/hmg/ddr315 (2011).

16. Reddy, T. E. et al. Effects of sequence variation on differential allelic transcription factor occupancy and gene expression. Genome Res 22, 860–869, doi: 10.1101/gr.131201.111 (2012).

17. Forrest, A. R. et al. A promoter-level mammalian expression atlas. Nature 507, 462–470, doi: 10.1038/nature13182 (2014).

18. Andersson, R. et al. An atlas of active enhancers across human cell types and tissues. Nature 507, 455–461, doi: 10.1038/nature12787 (2014).

19. Akbani, R. et al. A pan-cancer proteomic perspective on The Cancer Genome Atlas. Nat Commun 5, 3887, doi: 10.1038/ncomms4887 (2014).

20. Rosenbloom, K. R. et al. ENCODE whole-genome data in the UCSC Genome Browser: update 2012. Nucleic Acids Res 40, D912–917, doi: gkr1012[pii]10.1093/nar/gkr1012 (2012).

21. Cotton, A. M. et al. Spread of X-chromosome inactivation into autosomal sequences: role for DNA elements, chromatin features and chromosomal domains. Hum Mol Genet 23, 1211–1223, doi: 10.1093/hmg/ddt513 (2014).

22. Marks, H. et al. Dynamics of gene silencing during X inactivation using allele-specific RNA-seq. Genome Biol 16, 149, doi: 10.1186/s13059-015-0698-x (2015).

23. Arner, E. et al. Gene regulation. Transcribed enhancers lead waves of coordinated transcription in transitioning mammalian cells. Science 347, 1010–1014, doi: 10.1126/science. 1259418 (2015).

24. Breiman, L. Random Forests. Machine learning 45, 5–32 (2001).

25. Sirchia, S. M. et al. Loss of the inactive X chromosome and replication of the active X in BRCA1-defective and wild-type breast cancer cells. Cancer Res 65, 2139–2146, doi: 10.1158/0008-5472.CAN-04-3465 (2005).

26. Kawakami, T. et al. The roles of supernumerical X chromosomes and XIST expression in testicular germ cell tumors. J Urol 169, 1546–1552, doi: 10.1097/01.ju.0000044927.23323.5a (2003).

27. Carrel, L. & Willard, H. F. X-inactivation profile reveals extensive variability in X-linked gene expression in females. Nature 434, 400–404, doi: 10.1038/nature03479 (2005).

28. Weinstein, J. N. et al. The Cancer Genome Atlas Pan-Cancer analysis project. Nat Genet 45, 1113–1120, doi: 10.1038/ng.2764 (2013).

29. Mathelier, A. et al. JASPAR 2014: an extensively expanded and updated open-access database of transcription factor binding profiles. Nucleic Acids Res 42, D142–147, doi: 10.1093/nar/gkt997 (2014).

30. Usheva, A. & Shenk, T. TATA-binding protein-independent initiation: YY1, TFIIB, and RNA polymerase II direct basal transcription on supercoiled template DNA. Cell 76, 1115–1121 (1994).

31. Cusanovich, D. A., Pavlovic, B., Pritchard, J. K. & Gilad, Y. The functional consequences of variation in transcription factor binding. PLoS Genet 10, e1004226, doi: 10.1371/journal.pgen,1004226 (2014).

32. Sigova, A. A. et al. Transcription factor trapping by RNA in gene regulatory elements. Science 350, 978–981, doi: 10.1126/science.aad3346 (2015).

33. Thorvaldsen, J. L., Weaver, J. R. & Bartolomei, M. S. A YY1 bridge for X inactivation. Cell 146, 11–13, doi: 10.1016/j.cell.2011.06.029 (2011).

34. Atchison, M. L. Function of YY1 in Long-Distance DNA Interactions. Front Immunol 5, 45, doi: 10.3389/fimmu.2014.00045 (2014).

35. Moore, B. L., Aitken, S. & Semple, C. A. Integrative modeling reveals the principles of multi-scale chromatin boundary formation in human nuclear organization. Genome Biol 16, 110, doi: 10.1186/s13059-015-0661-x (2015).

36. Schwalie, P. C. et al. Co-binding by YY1 identifies the transcriptionally active, highly conserved set of CTCF-bound regions in primate genomes. Genome Biol 14, R148, doi: 10.1186/gb-2013-14-12-r148 (2013).

37. Minajigi, A. et al. Chromosomes. A comprehensive Xist interactome reveals cohesin repulsion and an RNA-directed chromosome conformation. Science 349, doi: 10.1126/science.aab2276 (2015).

38. Heard, E. & Bickmore, W. The ins and outs of gene regulation and chromosome territory organisation. Curr Opin Cell Biol 19, 311–316, doi: 10.1016/j.ceb.2007.04.016 (2007).

39. Deng, X. et al. Bipartite structure of the inactive mouse X chromosome. Genome Biol 16, 152, doi: 10.1186/s13059-015-0728-8 (2015).

40. Rahl, P. B. et al. c-Myc regulates transcriptional pause release. Cell 141, 432–445, doi: 10.1016/j.ce11.2010.03.030 (2010).

41. Adelman, K. & Lis, J. T. Promoter-proximal pausing of RNA polymerase II: emerging roles in metazoans. Nat Rev Genet 13, 720–731, doi: 10.1038/nrg3293 (2012).

42. Velia, P., Barozzi, I., Cuomo, A., Bonaldi, T. & Pasini, D. Yin Yang 1 extends the Myc-related transcription factors network in embryonic stem cells. Nucleic Acids Res 40, 3403–3418, doi: 10.1093/nar/gkr1290 (2012).

43. Gordon, S., Akopyan, G., Garban, H. & Bonavida, B. Transcription factor YY1: structure, function, and therapeutic implications in cancer biology. Oncogene 25, 1125–1142, doi: 10.1038/sj.onc.1209080 (2006).

44. Nilsson, J. A. & Cleveland, J. L. Myc pathways provoking cell suicide and cancer. Oncogene 22, 9007–9021, doi: 10.1038/sj.onc.1207261 (2003).

45. Chaligne, R. et al. The inactive X chromosome is epigenetically unstable and transcriptionally labile in breast cancer. Genome Res 25, 488–503, doi: 10.1101/gr.185926.114 (2015).

46. Team, R. C. R: A Language and Environment for Statistical Computing, <http://www.R-proiect.org/> (2015).

47. Huber, W. et al. Orchestrating high-throughput genomic analysis with Bioconductor. Nat Methods 12, 115–121, doi: 10.1038/nmeth.3252 (2015).

48. Cunningham, F. et al. Ensembl 2015. Nucleic Acids Res 43, D662–669, doi: 10.1093/nar/gku1010 (2015).

49. Wiener, A. L. a. M. Classification and Regression by randomForest. R News 2, 18–22 (2002). <http://CRAN.R-proiect.org/doc/Rnews/>.

50. Chen, H. & Boutros, P. C. VennDiagram: a package for the generation of highly-customizable Venn and Euler diagrams in R. BMC Bioinformatics 12, 35, doi: 10.1186/1471-2105-12-35 (2011).

51. Fortin, J. P. et al. Functional normalization of 450k methylation array data improves replication in large cancer studies. Genome Biol 15, 503, doi: 10.1186/s13059-014-0503-2 (2014).

52. Mathelier, A. et al. JASPAR 2016: a major expansion and update of the open-access database of transcription factor binding profiles. Nucleic Acids Res, doi: 10.1093/nar/gkv1176 (2015).

53. Pohl, A. & Beato, M. bwtool: a tool for bigWig files. Bioinformatics 30, 1618–1619, doi: 10.1093/bioinformatics/btu056 (2014).

54. Abecasis, G. R. et al. An integrated map of genetic variation from 1,092 human genomes. Nature 491, 56–65, doi: 10.1038/nature11632 (2012).

